# A standard area diagram for potato common scab: comparable performance of image- and object-based validation

**DOI:** 10.64898/2026.03.18.712681

**Authors:** Luis Ignacio Cazón, Juan Andrés Paredes, María Quiroga, Fabiana Guzmán

## Abstract

Potato common scab (*Streptomyces* sp.) is an economically important disease that reduces the quality and market value of tubers. A key aspect in developing management strategies involves accurately quantifying the disease. Due to the three-dimensional nature of the tuber and the heterogeneous distribution of lesions across its surface, visual estimates of severity can be challenging. Therefore, the objectives of this study were to develop and validate a standard area diagram (SAD) for estimating common scab severity on potato tubers and to compare validation outcomes obtained using real tubers and digital images. A SAD comprising six severity levels (from 1.3 to 66.8%) was developed based on image analysis of naturally infected tubers. Validation was conducted using two complementary approaches in which inexperienced raters evaluated either real potato tubers or digital images of the same tubers under unaided and aided conditions. Accuracy, bias components, and inter-rater reliability were quantified using absolute error metrics, Lin’s concordance correlation coefficient, intraclass correlation coefficients, and overall concordance correlation coefficients. Use of the SAD significantly improved accuracy, reduced systematic bias, and increased inter-rater reliability across both validation approaches. No significant differences were detected between assessments conducted on real tubers and images, although image-based evaluations showed a slight, non-significant tendency toward reduced scale and location bias under aided conditions. These results demonstrate that a dimension-aware SAD integrating information across the full tuber surface enhances the reliability and reproducibility of visual severity assessments and supports the use of image-based evaluations for training, large-scale surveys, and remote or collaborative applications involving three-dimensional plant organs.

## INTRODUCTION

Potato (*Solanum tuberosum* L.) is one of the most important food crops worldwide. According to FAO statistics, global potato production reached approximately 383 million tons in 2023, ranking fifth after sugarcane, maize, rice, and wheat, and playing a key role in sustaining global food security (Batool et al. 2020; Tao et al. 2023; FAOSTAT 2024). Potato production is geographically concentrated, with a limited number of countries accounting for most of the global output. China and India together contribute nearly one-third of total production, followed by Ukraine, the United States, and Russia (FAOSTAT 2024; Mishra et al. 2024). Despite this concentration in production, potatoes can be cultivated in diverse agroecological regions, encompassing temperate, subtropical, and high-altitude environments. This geographic breadth is associated with marked differences in cropping systems, seed sources, management practices, and phytosanitary pressures. Such variability contributes to heterogeneous disease dynamics and management challenges across production areas (INTA 2023).

From a phytosanitary perspective, common scab of potato, caused by *Streptomyces* spp., is among the most economically important diseases affecting the crop. Common scab is a monocyclic disease characterized by corky lesions on tubers that reduce quality and directly affect market value (Tao et al. 2023). Several species within the genus Streptomyces are associated with the disease, with *Streptomyces scabies* considered the most important. Management strategies include crop rotation, soil pH adjustment, irrigation management, and the use of less susceptible cultivars (Li et al. 2021; Porto et al. 2025). However, due to the diversity of species involved in the common scab complex and their differential responses to environmental conditions, disease intensity can be highly variable across production regions, resulting in substantial economic losses worldwide (INTA 2023).

Accurate disease quantification is a fundamental component for assessing the effectiveness of management strategies and for epidemiological and applied research purposes (Bock et al. 2015; Del Ponte 2024; Cazón et al. 2025; Coronel et al. 2025). Among the parameters commonly used, disease severity, defined as the proportion of the plant organ surface affected by disease, is particularly relevant (Bock et al. 2021). In the case of common scab, severity is a critical variable because tuber quality and commercial value are directly determined by the proportion of surface area affected by lesions.

Despite the development of advanced technologies for disease quantification, visual estimation remains widely used in plant pathology (Bock et al. 2022). Visual assessment consists of assigning a value (e.g., a score or percentage) to the severity of symptoms perceived by the human eye (Bock et al. 2021). This approach is time- and cost-efficient, adaptable to different workloads, and does not require specialized infrastructure or expensive equipment. However, visual estimation also presents limitations, including difficulties in evaluating large areas and errors associated with evaluator subjectivity and experience (Bock et al. 2010, 2020; Del Ponte et al. 2021; Cazón et al. 2025)

In addition to evaluator-related factors, symptom patterns and the characteristics of the affected organ influence the accuracy of visual assessments. Factors such as lesion size, shape, number, and degree of coalescence, as well as the dimensionality of the affected organ, can substantially influence estimation accuracy (Del Ponte et al. 2021). Estimating disease severity on leaves, which are essentially two-dimensional structures, differs markedly from assessments on stems, fruits, or tubers, which are three-dimensional organs. Among the tools developed to reduce errors associated with these factors are standard area diagrams (SADs) (Del Ponte et al. 2017, 2021; Bock et al. 2020, 2022). SADs are pictorial or graphic representations—typically photographs or drawings—illustrating increasing levels of disease severity on plant organs, usually expressed as percentages to facilitate interpolation (Del Ponte et al. 2017; Bock et al. 2021; Cazón et al. 2025). Validation of SADs is commonly performed using images that represent different severity levels on the affected organ. However, when dealing with three-dimensional structures such as potato tubers, image-based validation may introduce biases if the images do not adequately represent the entire surface area of the organ. Additional sources of bias may arise from image standardization procedures, including uniformity in color, size, and shape, which may differ from the natural variability observed in real tubers under field or commercial conditions.

Therefore, the objectives of this study were to (i) develop and validate a SAD for estimating the severity of common scab on potato tubers, ensuring full surface representation in each diagram, and (ii) determine whether differences exist between validation processes conducted using real tubers and digital images.

## MATERIALS AND METHODS

### SAD development and design

A total of 135 potato tubers exhibiting different levels of common scab severity were collected in March 2025 from commercial production fields located in the Villa Dolores region, Córdoba, Argentina (31°55′23.3”S, 65°26′32.1”W). Each tuber was cut longitudinally in half using a knife, and both external surfaces were digitized in a single image on a white background using a smartphone camera. Images were stored in JPG format at a resolution of 200 dpi.

The images were processed using *Adobe Photoshop* software (Adobe Systems Inc., 2011) to remove small shadows generated by tuber edges on the white background that could interfere with severity quantification. The percentage of area affected by *Streptomyces* sp. was determined using the *pliman* package (Olivoto, 2022) in *R* (R Core Team, 2022), which performs automated image analysis through segmentation methods to differentiate diseased from healthy tissue. Before image processing with *pliman*, lesions in each image were manually colored red to facilitate software-based segmentation (Fig. 1).

**Figure 1.**
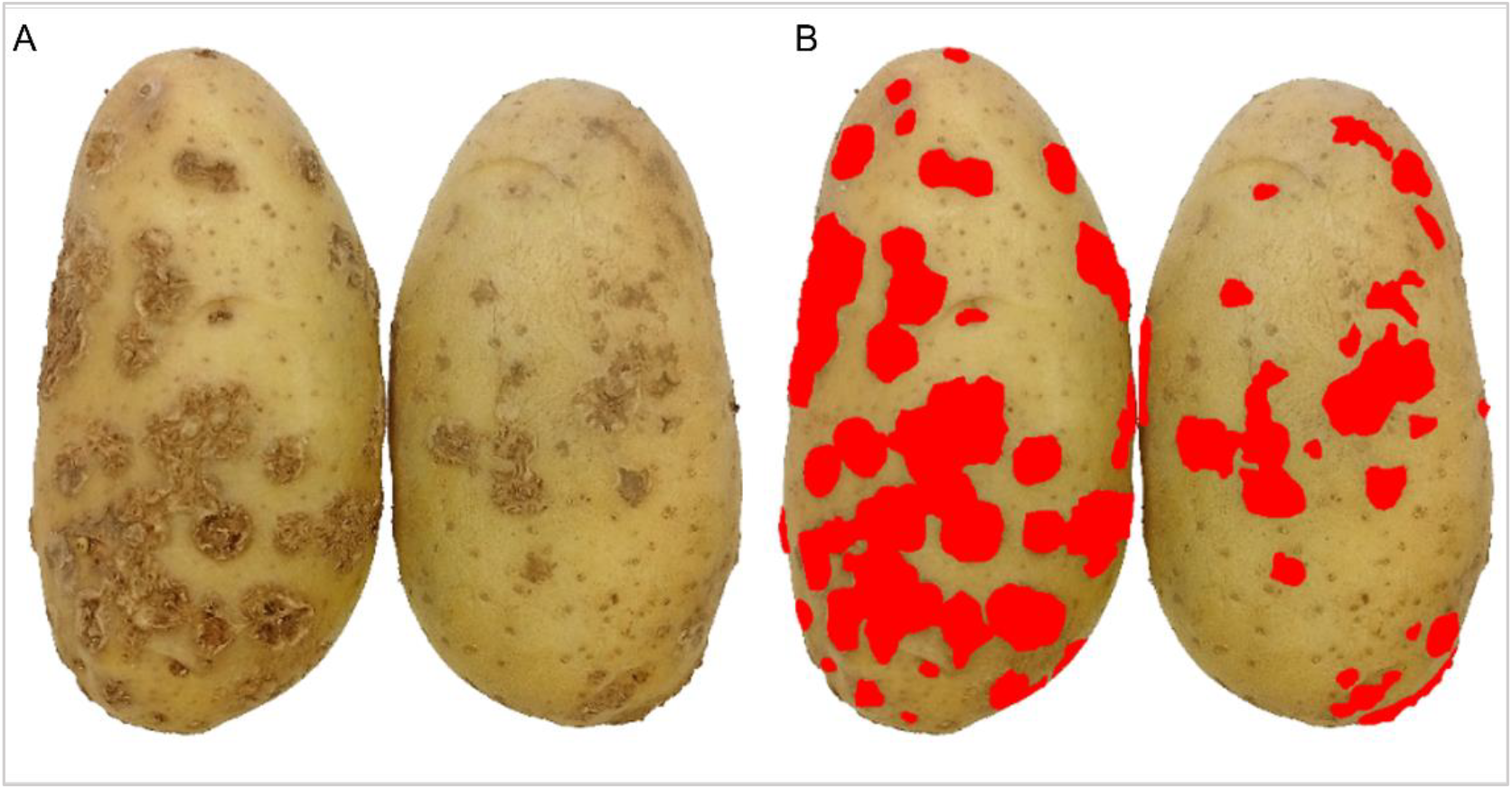
(A) Longitudinally cut potato tuber showing both exposed surfaces with symptoms of common scab. (B) The same tuber, with scab lesions digitally highlighted.

The SAD was designed using *Microsoft PowerPoint* (Microsoft, 2024) based on the minimum and maximum severity levels observed in the sampled tubers. Diagrams were distributed at linear intervals, and an additional diagram at low severity levels was included to reduce overestimation at the lower end of the scale, as recommended by Liu et al. (2019).

### SAD validation

Two complementary validation approaches were used: validation with real potato tubers and validation with potato images.

#### Validation with real potato tubers

Eighty potato tubers exhibiting common scab symptoms were harvested in August 2025 from commercial fields in the Villa Dolores region, Córdoba, Argentina. From this group, 40 tubers representing the full range of observed severity levels were selected. Each tuber was identified with a unique number (1 to 40) to ensure traceability throughout the validation process. Validation was conducted in two consecutive stages by 21 raters with no prior experience in potato disease assessment. In the first stage, raters estimated disease severity without the aid of the SAD (unaided evaluation). In the second stage, severity was estimated using the SAD (aided evaluation). In both stages, a fixed evaluation time of 20 seconds per tuber was imposed.

#### Validation with potato images

For the second approach, the same 40 tubers used in the real-potato validation were digitized (maintaining their original number identity) using the same procedure described above for SAD development. To determine the actual severity of common scab on these tubers, lesions were manually colored using *Microsoft PowerPoint* (Microsoft, 2024), and the images were processed with *pliman*. The resulting severity values were used as reference standards for both validation approaches. All 40 images were compiled into a PDF file and distributed to 18 raters with no prior experience in potato disease assessment. The validation was performed in two stages (unaided and aided). As in the real-potato validation, a fixed evaluation time of 20 seconds per image was applied.

### Accuracy and reliability

Accuracy and reliability were assessed by calculating the absolute error (difference between estimated and actual severity) and its standard deviation, which was used as an indicator of error variability, for each rater and assessment round (Del Ponte et al., 2023). Agreement between actual and estimated severity was quantified using Lin’s concordance correlation coefficient (*CCC*) (Lin, 1989; Madden et al., 2007; Del Ponte et al., 2017).

Accuracy components included overall concordance (*ρc*), precision (Pearson’s *r*), generalized bias coefficient (*Cb*), scale bias (*v*), and location bias (*u*), as previously described (Dolinski et al., 2017; Coronel et al., 2025). These parameters were analyzed using generalized linear mixed-effects models, including the interaction between evaluation aid (unaided vs. aided) and validation approach (real potato tubers vs. potato images) as fixed effects, and raters as random intercepts. Model assumptions were verified using the *DHARMa R* package based on residual simulation (Hartig, 2021), and mean comparisons were performed using Tukey’s test.

### Inter-rater reliability

Inter-rater reliability was assessed using the intraclass correlation coefficient (*ICC*) based on the 95% confidence intervals (Shoukri and Pause 1999) and the overall concordance correlation coefficient (*OCCC*), an extension of Lin’s CCC for multiple raters (Barnhart et al. 2002). As described by Barnhart et al. (2007), different forms of ICC can be defined depending on the study design, with estimates obtained from appropriate variance components (Shrout & Fleiss, 1979) when no reference evaluator is defined. In this study, *ICC2* was calculated using the *ICC* function of the *psych R* package (with the *lmer* argument set to TRUE) to obtain both point estimates and confidence intervals, following the approach of Pereira et al. (2020). The *OCCC* was calculated using the *epi*.*occc* function of the *epiR R* package (Stevenson et al., 2019).

## RESULTS

### SAD development and design

The proposed standard area diagram (SAD) comprises six true-color images representing common scab severity levels ranging from 1.3 to 66.8% (Fig. 2). Each diagram depicts a potato tuber cut longitudinally in half, displaying both external surfaces. Severity classes were defined using intervals of approximately 15% between consecutive diagrams. In addition, an intermediate diagram corresponding to 9.9% severity was included between the minimum class and the 21.9% severity level to reduce overestimation at low severity values, resulting in an amended linear percentage scale.

**Figure 2.**
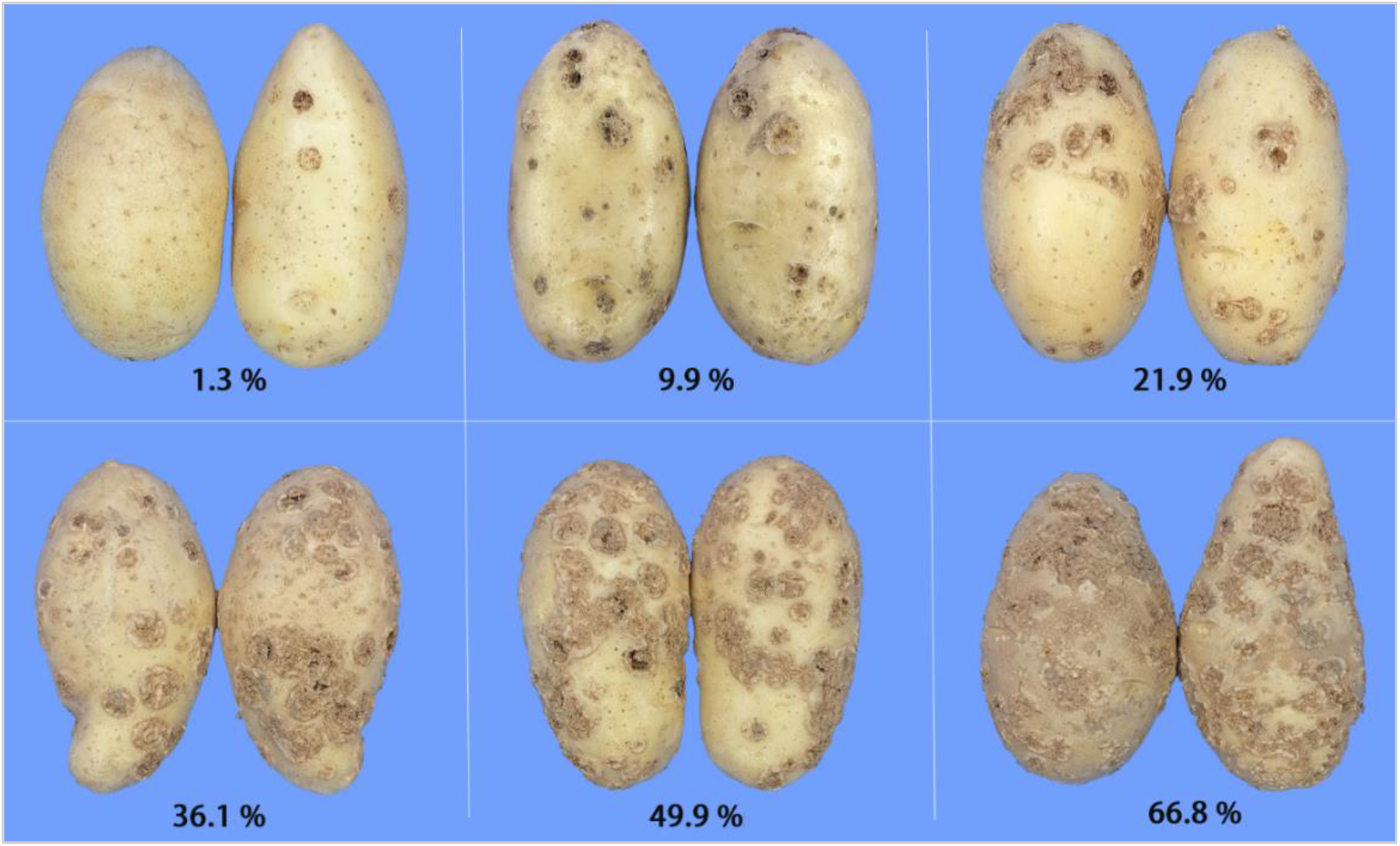
Standard area diagrams (SAD) set for visual estimation of the severity of Potato common scab, caused by *Streptomyces* sp.

### SAD validation

The use of the SAD significantly reduced both the magnitude and variability of severity estimation errors across the two validation approaches (P < 0.05; Fig. 3A, B). Specifically, the standard deviation of the estimation error decreased by 8.09 units (from 24.92 to 16.83) in the validation with real potato tubers, and by 7.63 units (from 23.68 to 16.05) in the image-based validation (Table 1), indicating improved inter-rater reliability (Figure 3A, B).

**Table 1.** Effect of the SAD set to improve the visual severity estimation of Potato common scab on the LCCC (*ρ*_*c*_), precision (*r*), bias coefficient (*C*_*b*_), location bias (*u*), scale bias (*v*), standard deviation (*Sd*), agreement between raters described by the intraclass correlation coefficient (ICC) at 95% of the confidence interval (95% CI). Different letters within the same column for each approach indicate significant difference (Tukey’s test, P ≤ 0.05)

| | | $\rho_c$ | $r$ | $C_b$ | $Sd$ | $u$ | $v$ | ICC (95% CI) |
| --- | --- | --- | --- | --- | --- | --- | --- | --- |
| Real Potatoes | Unaided | 0.68a | 0.87a | 0.79a | 24.92 | 2.49 | 1.30 | 0.57 |
|  | Aided | 0.87b | 0.90b | 0.96b | 16.83 | 1.50 | 1.11 | 0.85 |
| Images Potatoes | Unaided | 0.65a | 0.86a | 0.80a | 23.68 | 4.97 | 1.22 | 0.49 |
|  | Aided | 0.84b | 0.89a | 0.96b | 16.05 | -0.90 | 1.04 | 0.77 |
| Global process | Unaided | 0.66a | 0.87a | 0.80a | 24.09 | 3.63 | 1.26 | 0.53 |
|  | Aided | 0.85b | 0.90b | 0.96b | 16.52 | 0.40 | 1.08 | 0.81 |

**Figure 3.**
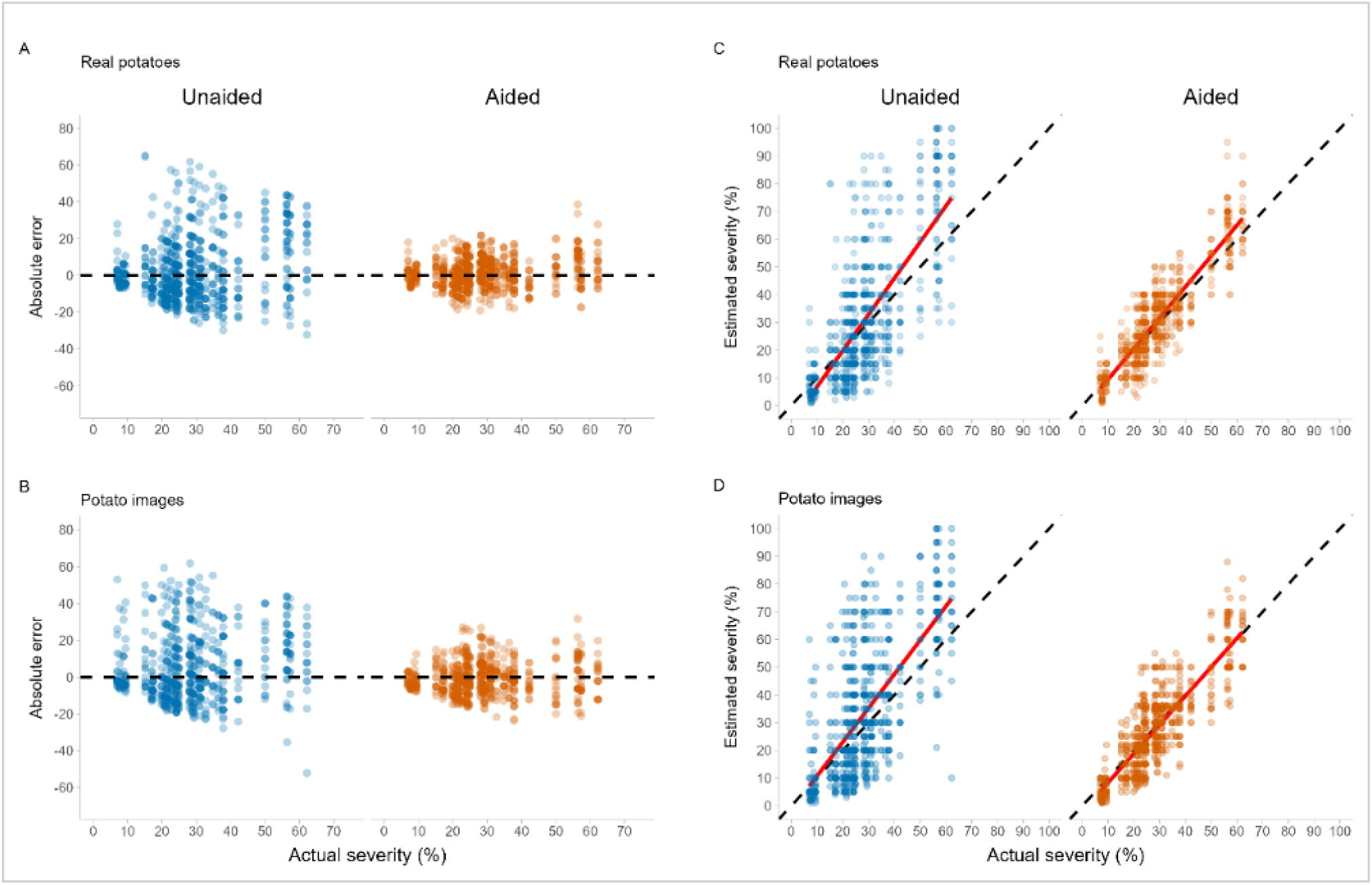
Performance of severity estimates obtained with and without the standard area diagram (SAD) of potato common scab using two validation approaches (real potatoes and potato images). Panels A and B show the absolute error of severity estimates as a function of actual severity for real potato tubers (A) and potato images (B), under unaided and aided conditions. Panels C and D show the relationship between estimated and actual severity for real potato tubers (C) and potato images (D). Dashed black lines indicate the 1:1 line (perfect agreement), and solid red lines represent the fitted regression lines. Blue and orange points correspond to unaided and aided estimates, respectively.

This reduction in absolute error was accompanied by a marked decrease in systematic bias components. Location bias (*u*) was substantially reduced following the use of the SAD in both validation approaches, decreasing from 2.49 to 1.50 in the assessment conducted with real potato tubers and from 4.97 to −0.90 in the image-based validation (Table 1). A similar trend was observed in the global analysis, where *u* declined from 3.63 to 0.40. These improvements are visually supported by the closer alignment of regression lines with the identity line in Fig. 3C, D.

Similarly, scale bias (*v*) decreased consistently with the use of the SAD, approaching the ideal value of 1. In the real-potato validation, *v* declined from 1.30 to 1.11, while in the image-based validation it decreased from 1.22 to 1.04. This improvement is reflected in Fig. 3C and 3D by the reduction in slope deviations and the tighter clustering of estimates around the identity line under aided conditions. Considering the global process, scale bias was reduced from 1.26 to 1.08 (Table 1).

The SAD significantly improved all parameters associated with the accuracy and precision of severity estimates in both validation approaches, using real potato tubers and potato images. No significant differences were detected between validation approaches within the same evaluation condition (unaided vs. aided), indicating comparable performance of the SAD when applied to real objects and digital images (Table 1). Lin’s concordance correlation coefficient (CCC) increased by 0.19 in both validation approaches (Figure 4), as well as in the overall analysis following the use of the scale (Table 1).

**Figure 4.**
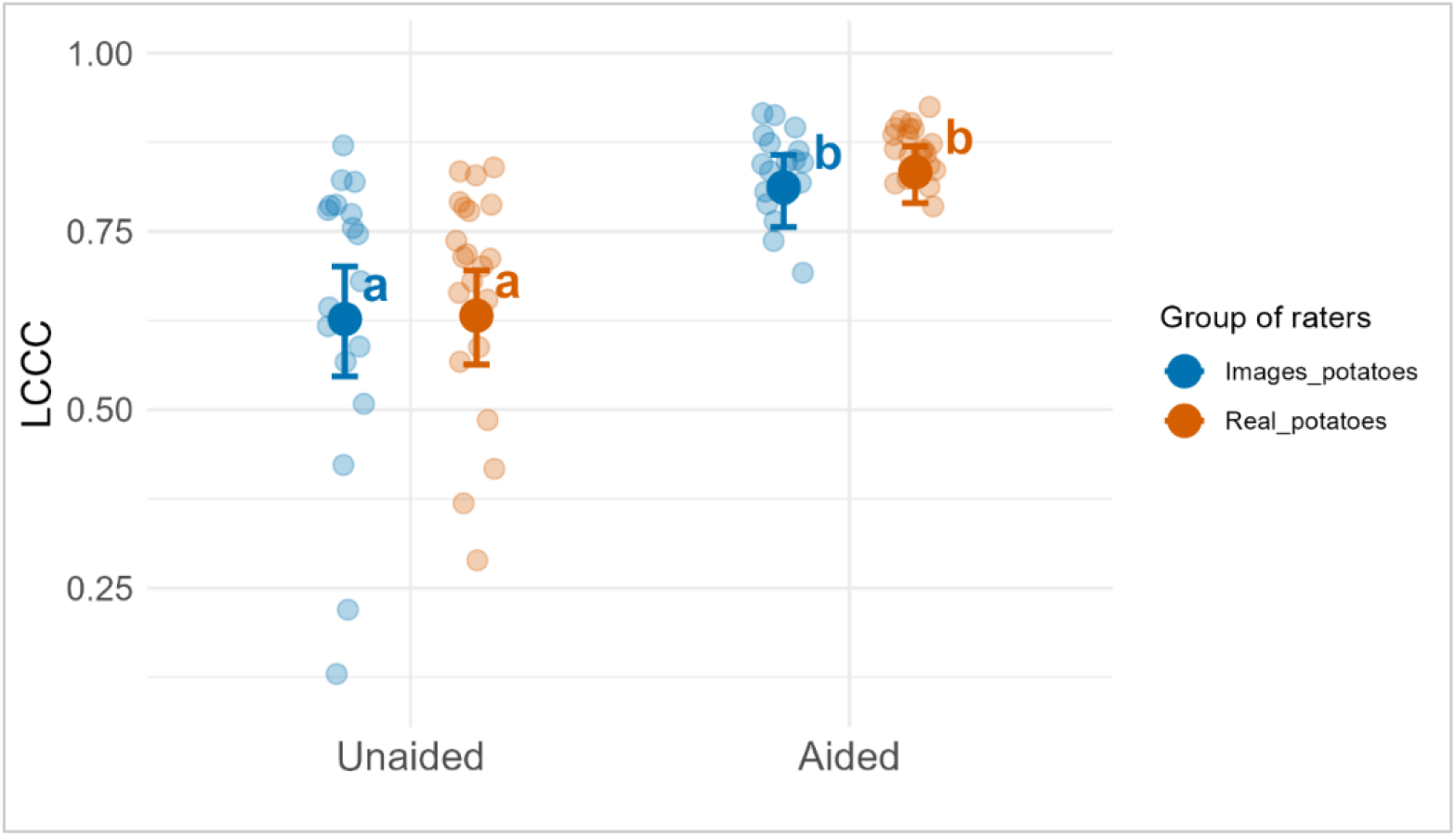
Lin’s concordance correlation coefficient (LCCC) for severity estimates obtained under unaided and aided conditions using two validation approaches. Points represent individual raters, while larger symbols with error bars indicate the mean LCCC ± standard error for each group. Blue symbols correspond to evaluations based on potato images, and orange symbols to evaluations using real potato tubers. Different letters indicate significant differences between conditions (P < 0.05).

Likewise, precision, expressed as Pearson’s correlation coefficient (r), increased significantly by 0.03 across all validation scenarios. All raters participating in the validation process exhibited gains in accuracy when using the SAD, regardless of the validation approach employed (Fig. 5).

**Figure 5.**
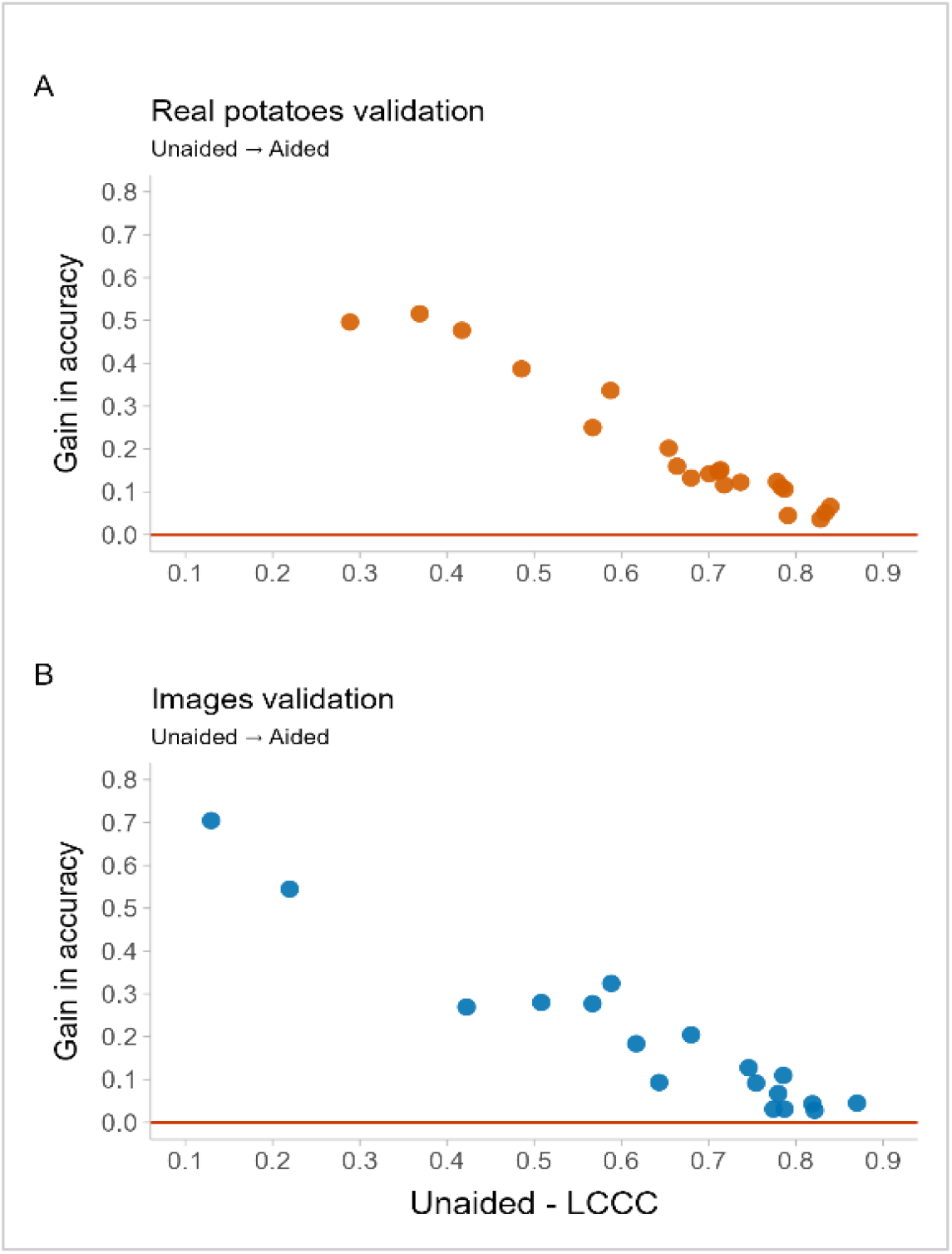
Gain in accuracy achieved by using the SAD as a function of baseline performance under unaided conditions. Panel A shows results from the real potato validation, and panel B from the image-based validation. Each point represents an individual rater. Gain in accuracy was calculated as the difference in Lin’s concordance correlation coefficient (LCCC) between aided and unaided assessments, while the x-axis shows the LCCC obtained under unaided conditions. The horizontal line at zero indicates no gain in accuracy.

Finally, both the intraclass correlation coefficient (ICC) and the overall concordance coefficient (OCC) increased substantially (by 0.28) in all validation approaches after the use of the SAD, indicating reduced inter-rater variability and enhanced agreement in disease severity assessments. In this study, the values for both parameters were similar; therefore, only the ICC values are presented in Table 1.

## DISCUSSION

Despite advances in AI-based technologies for disease quantification, visual estimation remains a valid and widely used approach for disease assessment in many pathosystems (Bock et al., 2010, 2020; Del Ponte et al., 2021). In this context, improving accuracy and reproducibility is essential to ensure reliable severity estimates. As highlighted by Del Ponte et al. (2021), visual estimation accuracy is influenced not only by evaluator-related factors but also by symptom attributes, such as lesion size, shape, number, and degree of coalescence, as well as by the dimensionality of the affected organ. Whereas severity assessments on leaves, which are essentially two-dimensional structures, require specific visual skills, evaluations on three-dimensional organs such as stems, fruits, and tubers pose additional cognitive and methodological challenges. In these cases, evaluators must integrate information across multiple surfaces, often relying on partial visual cues or mental extrapolation, which increases subjectivity and systematic error (Bock et al. 2010)

In the context of potato common scab, these limitations are particularly relevant because lesions are frequently irregular in shape, heterogeneous in size, and unevenly distributed over the tuber surface (Dees and Wanner 2012). Consequently, conventional visual assessments based on partial surface observation may introduce errors to accurately represent true disease severity. The results of the present study demonstrate that the use of a SAD explicitly designed to account for the three-dimensional nature of potato tubers can substantially mitigate these limitations. By providing a visual reference that integrates information from the entire tuber surface, the proposed SAD improved estimation accuracy and consistency across raters, reinforcing the value of organ-specific diagrammatic tools for disease quantification on three-dimensional plant structures.

The consistent reduction in absolute error and error variability observed when the SAD was used highlights the importance of visual aids for standardizing disease assessments, particularly among inexperienced raters (Del Ponte et al. 2017). These improvements were accompanied by marked reductions in both location bias (*u*) and scale bias (*v*), indicating that the SAD effectively corrected systematic tendencies toward overestimation at low severity levels and slope distortions across the severity range. Together, these results confirm that the proposed SAD not only improves accuracy but also enhances the biological interpretability of severity estimates.

A key contribution of this study is the explicit consideration of the limitations associated with evaluating three-dimensional plant organs using two-dimensional visual references. Although, to our knowledge, no diagrammatic scales have explicitly addressed organ three-dimensionality, studies in plant phenotyping have highlighted the importance of capturing the entire surface of plant organs to avoid the loss of spatial information and biased measurements inherent to two-dimensional assessments (Paulus, 2019). Potato tubers exhibit complex geometries, and the spatial distribution of common scab lesions often varies markedly across the tuber surface. By representing both external surfaces of a longitudinally cut tuber within each diagram, the proposed SAD provides a more comprehensive visual reference that better approximates the true disease burden. This design represents an advantage over the SAD proposed by Andrade et al. (2019), in which only a single tuber surface is depicted in each diagram. Although their SAD resulted in improved estimation accuracy, reliance on a single visible surface may limit its applicability when disease distribution is asymmetric or when evaluators are required to infer severity in unobserved areas. In contrast, the dual-surface representation adopted in the present study reduces the need for mental extrapolation, thereby lowering cognitive load and minimizing subjective bias during severity estimation (Bock et al. 2010).

An additional strength of this study is the use of two complementary validation approaches, real potato tubers and digital images, which allowed us to assess whether the performance of the proposed SAD was influenced by the evaluation methodology. Overall, no significant differences were detected in *ρ*_*c*_, *r*, and *Cb* between validation approaches under either unaided or aided conditions, indicating that the SAD performs consistently when applied to real three-dimensional objects and to two-dimensional image representations. This result suggests that, when properly designed and validated, image-based assessments can provide reliability levels comparable to those obtained from direct evaluations of physical plant organs. Although no statistically significant differences were observed between approaches, a slight visual tendency toward better fit was observed for image-based validation when using the SAD. Under these conditions, the regression line in Fig. 3D shows a closer alignment to the 1:1 identity line compared with the corresponding relationship observed for real tubers (Fig. 3C). This tendency is consistent with the observed reductions in both location bias (*u*) and scale bias (*v*) in the image-based assessments (Table 1). The lower values of *u* indicate a decrease in systematic over- or underestimation across the severity range, while *v* values closer to unity reflect improved proportional agreement between estimated and actual severity (Dolinski et al., 2017, Coronel et al., 2025). Together, these results suggest that the use of standardized images may slightly facilitate bias correction when aided by the SAD, likely by reducing visual complexity and aiding evaluators in calibrating severity estimates. Nevertheless, given the absence of statistically significant differences between validation approaches, this pattern should be interpreted cautiously and must be explored in future studies.

Additionally, the observed increases in inter-rater reliability (ICC and OCCC values) indicate that the SAD promotes greater consistency among evaluators, an essential requirement for epidemiological studies, cultivar screening, and fungicide or management trials (Nutter et al., 1993, 2006; Madden et al., 2007). The similarity between ICC and OCCC values further suggests that the scale performs robustly regardless of the statistical framework used to quantify agreement (Lin 1989). The convergence of these two metrics indicates that the use of the SAD not only reduces random variability of each rater but also limits systematic disagreement between raters (Shoukri and Pause, 1999; Barnhart et al., 2002). This indicator reinforces the suitability of the proposed SAD scale for applications requiring reproducible severity assessments across multiple raters and assessment contexts.

From a practical standpoint, the results of this study highlight the usefulness of the proposed SAD as a flexible tool applicable across different assessment contexts. The comparable performance observed between assessments conducted on real potato tubers and digital images supports the use of image-based evaluations for assessor training and for remote or collaborative studies in which access to physical samples may be limited. This approach is particularly relevant for online training platforms, such as TraineR2 (Del Ponte et al., 2023), where standardized image sets representing common scab symptoms can be integrated with validated SADs to enhance and harmonize assessment skills.

From a methodological perspective, the findings suggest that the choice between image-based and object-based assessments should be guided by the study objectives and logistical constraints, rather than by concerns about the effect of pathosystem characteristics on raters, provided that the SAD adequately represents the three-dimensional nature of the organ being assessed.

## CONCLUSION

This study presents the development and validation of a standard area diagram (SAD) for the visual estimation of common scab severity on potato tubers, explicitly designed to account for the three-dimensional nature of the affected organ. The proposed SAD significantly improved the accuracy, precision, and reproducibility of severity estimates, reducing both systematic and random evaluator-related errors across different assessment conditions.

Importantly, the comparable performance observed between validations conducted using real potato tubers and digital images demonstrates the robustness and versatility of the scale. These findings support the use of image-based assessments as a reliable alternative for evaluator training, provided that the diagrammatic references adequately represent the full surface complexity of the organ being assessed.

The approach described in this study may be extended to other pathosystems involving three-dimensional plant organs, contributing to more standardized and comparable disease quantification in both research and applied contexts.

## Acknowledgements

We wish to thank INTA for providing resources for compiling this project. We also want to thank the volunteers who participated in the SAD validation process and potato producers from Villa Dolores (Córdoba – Argentina) who collaborated with the field material.

## Author’s contribution

**LIC** and **JAP** conceptualized the study, wrote the manuscript, contributed to the SAD set development, validation process, and data analysis. **MQ** contributed to the SAD set validation process, image processing, and co-wrote the manuscript. **FG** conceptualized the study, collected the samples, and contributed to the SAD set development and validation.

## Data Availability

The datasets generated and/or analyzed during the current study are available at the following repository link: (https://repositorio.inta.gob.ar/xmlui/handle/20.500.12123/25508)

